# CRISPR/Cas9 gene editing for the creation of an MGAT1 deficient CHO cell line to control HIV-1 vaccine glycosylation

**DOI:** 10.1101/368357

**Authors:** Gabriel Byrne, Sara M. O’Rourke, David L. Alexander, Bin Yu, Rachel C. Doran, Meredith Wright, Qiushi Chen, Parastoo Azadi, Phillip W. Berman

**Affiliations:** Department of Biomolecular Engineering, University of California Santa Cruz, Santa Cruz, California United States of America.; Complex Carbohydrate Research Center, University of Georgia, Athens, Georgia, United States of America.

**Keywords:** HIV vaccine, glycosylation, CHO cells, neutralizing antibodies, gene editing, CRISPR/Cas9, gp120

## Abstract

Over the last decade multiple broadly neutralizing monoclonal antibodies (bN-mAbs) to the HIV-1 envelope protein, gp120, have been described. Surprisingly many of these recognize epitopes consisting of both amino acid and glycan residues. Moreover, the glycans required for binding of these bN-mAbs are early intermediates in the N-linked glycosylation pathway. This type of glycosylation substantially alters the mass and net charge of HIV envelope (Env) proteins compared to molecules with the same amino acid sequence but possessing mature, complex (sialic acid containing) carbohydrates. Since cell lines suitable for biopharmaceutical production that limit N-linked glycosylation to mannose-5 (Man_5_) or earlier intermediates are not readily available, the production of vaccine immunogens displaying these glycan dependent epitopes has been challenging. Here we report the development of a stable suspension adapted CHO cell line that limits glycosylation to Man_5_ and earlier intermediates. This cell line was created using the CRISPR/Cas9 gene editing system and contains a mutation that inactivates the gene encoding Mannosyl (Alpha-1,3-)-Glycoprotein Beta-1,2-N-Acetylglucosaminyltransferase (MGAT1). Monomeric gp120s produced in the MGAT1^-^ CHO cell line exhibit improved binding to prototypic glycan dependent bN-mAbs directed to the V1/V2 domain (e.g. PG9) and the V3 stem (e.g. PGT128 and 10–1074) while preserving the structure of the important glycan independent epitopes (e.g. VRC01). The ability of the MGAT1-CHO cell line to limit glycosylation to early intermediates in the N-linked glycosylation pathway, without impairing the doubling time or ability to grow at high cell densities, suggest that it will be a useful substrate for the biopharmaceutical production of HIV-1 vaccine immunogens.

## Introduction

Despite thirty years of research, a vaccine capable of providing protection against human immunodeficiency virus type 1 (HIV) has yet to be described. However, considerable progress towards this goal has been achieved with the elucidation of the 3-dimensional structure of the HIV-1 envelope proteins (monomeric gp120 and trimeric gp140) and the characterization of multiple broadly neutralizing monoclonal antibodies (bN-mAbs) (1–5). As headway toward a protective vaccine continues, the practicalities of large-scale vaccine production must be addressed. A growing body of evidence indicates that the N-linked glycosylation structure will be a critical factor in both the design and manufacture of any HIV vaccine (6–8).

Beginning in 2009, we learned that multiple bN-mAbs recognized glycan dependent epitopes on the HIV envelope protein, gp120. In an unanticipated development, several families of bN-mAbs require mannose-5 (Man_5_)and/or mannose-9 (Man_9_) for binding to key epitopes of gp120 (6, 9–11). As these bN-mAbs were being described, the data from the RV144 HIV vaccine trial was released. This study provided evidence for the first time that vaccination could prevent HIV infection in humans (12). The regimen used in this trial involved immunization with a bivalent gp120 vaccine (AIDSVAX B/E) to stimulate an antibody response as well as immunization with a recombinant canarypox vector to stimulate a cell mediated immune response (13–15). This immunization protocol resulted in modest (31.2%) but significant vaccine efficacy (12). Examination of the gp120 subunit vaccines used in the RV144 trial showed that both components (MN-rgp120 and A244-rgp120) were enriched for complex, sialic acid containing glycans and lacked the high-mannose glycosylation found on the surface of virions and native envelope proteins required to bind the new class of glycan dependent bN-mAbs(16–20). Thus, differences in glycosylation between the vaccine immunogens from the RV144 trial and virus particles could, in part, explain the low efficacy of RV144 and other gp120 based vaccines and their inability to elicit broadly neutralizing antibodies (bN-mAbs). Previously we reported that the same gp120s used in the RV144 trial could be modified to bind multiple bN-mAbs when expressed in a cell line (HEK 293 GnTI^-^) that limited N-linked glycosylation to Man_5_, or earlier species (e.g. Man_8_, Man_9_) (21). While in theory this cell line could be used to produce a glycan optimized gp120 vaccine, in reality this is not practical. The HEK 293 GnTI^-^ system is not suitable for clinical and large-scale production due to genetic instability and the inability to grow for sustained periods at high cell densities (22, 23).

CHO cells have long been the substrate of choice for the production of therapeutic glycoproteins. This is due to their ability to grow at high densities in serum-free suspension cultures, sustain high levels of protein expression over prolonged fermentation cycles, and incorporate complex glycans on exogenously expressed proteins (24–26). Typical glycoproteins contain only a few N-linked glycans, which aid in protein folding, intracellular trafficking. When these glycans terminate in sialic acid residues they increase resistance to proteolysis and extend serum half-life *in vivo*(27–29). Because of these physical and pharmacokinetic benefits, recombinant glycoprotein expression efforts have historically focused on maximizing the amount of complex, sialic acid containing glycans per molecule. Although modern production technology provides the means to express and purify properly folded recombinant glycoproteins at large scale, controlling the glycosylation has been a persistent problem for most glycoproteins due to the “non-templated” nature of glycosylation (30–32). The final glycan structure of proteins with only a few N-linked glycosylation sites can be highly variable with respect to the glycan structure branching, saccharides present, sialic acid content, and net charge. Glycosylation heterogeneity is known to result from a variety of variables including: cell type, protein expression levels, cell culture conditions, monosaccharide donor availability, and protein structure (30, 33–36). Controlling glycosylation heterogeneity in gp120 is particularly problematic due to the fact that it contains an average of 25 potential N-linked glycosylation sites (PNGS), comprising approximately 50% of the mass of the mature protein (37–40). Each glycan site may be different in composition than others on the same molecule or different at the same position from molecule to molecule. Variance is so great that 79 different glycan structures have been found to occur a single position in envelope proteins expressed in normal CHO cells (41).

In this paper we address the problems of glycosylation heterogeneity and bN-mAb binding in the large-scale production of recombinant envelope proteins by the development of a mutant CHO cell line (MGAT1^-^ CHO) in which the Mannosyl (Alpha-1,3-)-Glycoprotein Beta-1,2-N-Acetylglucosaminyltransferase (MGAT1) gene has been inactivated using CRISPR/Cas9 gene editing. The nomenclature for MGAT1 gene has changed over the years and was previously referred to as the GnTI gene. Inactivation or deficiency of the MGAT1 limits N-linked glycosylation to early oligo-mannose glycans (Man_5–9_) and enhances the binding of bN-mAbs to glycans dependent epitopes, as compared to earlier gp120 vaccines produced in normal CHO cells. Although other CHO cell lines, such as CHO Lec1, have been described that similarly limit oligomannose-structures, they grow slowly, and differ from parental cell lines in morphology and growth characteristics (42, 43). Thus, the development of a precision-engineered CHO cell line resulting from by CRISPR/Cas9 gene editing, is a desirable alternative for HIV vaccine manufacturing. This cell line should be useful for the production of stable cell lines suitable for the production HIV vaccines as well as other biopharmaceuticals where limiting the incorporation of sialic acid is beneficial.

## Results

### Silencing of CHO-S MGAT1 Gene

The goal of this project was to make an MGAT1 deficient CHO-S cell line using CRIPSR/Cas9. With this gene knocked out, complex, sialic acid containing, glycans cannot be formed and N-linked glycosylation is not processed beyond the oligomannose Man_5_ structure (Fig. 1). The CRISPR/Cas9 gene editing system allows for specific targeting of genes for deletion or modification by introducing double stranded breaks (DSB) followed by non-homologous end joining (NHEJ) or homology directed repair (HDR) (44, 45). We utilized a CRISPR/Cas9 nuclease vector containing an OFP reporter gene (Materials and Methods). After insertion of guide sequences, the vector contained all of the elements needed to induce a double stranded break in the MGAT1 gene. The sequence of the CHO MGAT1 gene was identified from GenBank, gene ID: 100682529 (46). Three target specific double stranded guide sequences were ligated into the vector between a U6 promoter and a tracrRNA sequence. The same vector encodes the Cas9 endonuclease and an orange fluorescent protein reporter gene, separated by a self-cleaving 2A peptide linker. This system allows for a single plasmid to encode for both the Cas9 and a complete gRNA, enabling the use of non-Cas9 expressing cells. Following ligation of these guide sequences, the vectors were transfected into CHO-S cells using the MaxCyte electroporation system (Fig. 2). Targets 1 and 2 were introduced individually; target 3 plasmid was mixed and added together in equal ratio with target 2, creating three separate pools of transfected cells. Twenty-four hours post transfection samples were serially diluted across five 96 well flat-bottoms plates at a calculated density of 0.5 cells per well. The plates were examined daily, and wells with more than a single colony was discarded. Across the 15 total plates, between 15 and 30 wells per plate contained single viable colonies that were transferred to 24 well plates upon reaching 20% confluency after 12 to 15 days. Wells in the 96 well plates that did not have at least several dozen cells by day 15 were discarded. A total of 166 colonies were expanded to 24 well plates: 55 from target 1 pool, 67 from target 2 pool, and 44 from combined target 2/3 pool.

**Fig 1.**
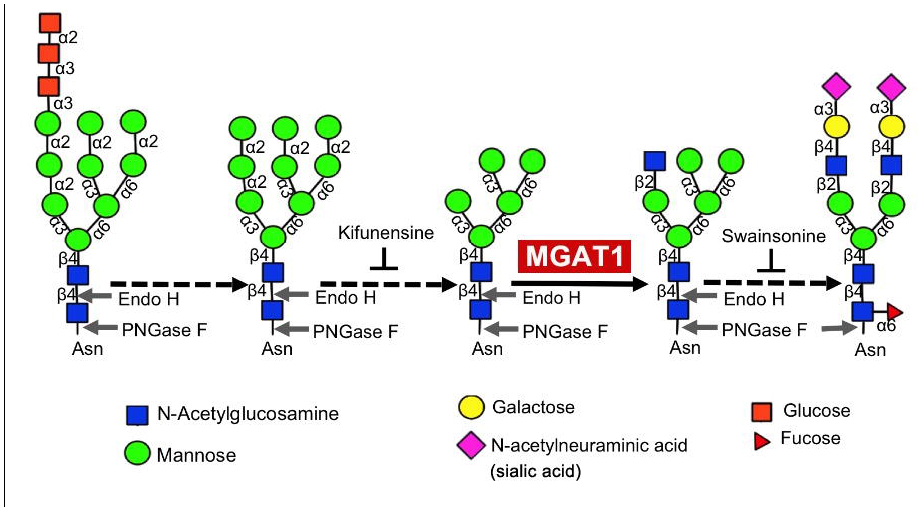
Simplified view of N-linked glycosylation pathway. N-linked glycosylation begins in the endoplasmic reticulum with the en-block transfer of a highly conserved Glc_3_Man_9_GlcNAc_2_ structure (left) to asparagine residues within the N-X-S/T motif of nascent proteins. This initial structure is sequentially trimmed to Man_9_GlcNAc_2_ and then Man_5_GlcNAc_2_ (center) as the protein moves from the ER to the Golgi apparatus. The enzyme, Mannosyl (Alpha-1,3-)-Glycoprotein (Beta-1,2)-N-Acetylglucosaminyltransferase (MGAT1, red box) adds an N-acetylglucosamine to the Man_5_ structure and is required to enable other glycosyltransferases to add monosaccharides creating hybrid (second from right) and complex (right) glycoforms. Treatment with endoglycosidase H (Endo H) cleaves simple, oligomannose containing glycans from glycoproteins, but not complex sialic acid containing glycans. PNGase F removes both simple and complex glycans from glycoproteins (indicated by the arrows). Kifunensine and swainsonine are inhibitors that halt processing at the steps indicated. Dashed black arrows indicate multiple enzymatic steps. Figure adapted from Binley, J.M., et al., *Role of Complex Carbohydrates in Human Immunodeficiency Virus Type 1 Infection and Resistance to Antibody Neutralization*. Journal of Virology, 2010. 84(11): p. 5637–5655. (47)

**Fig 2.**
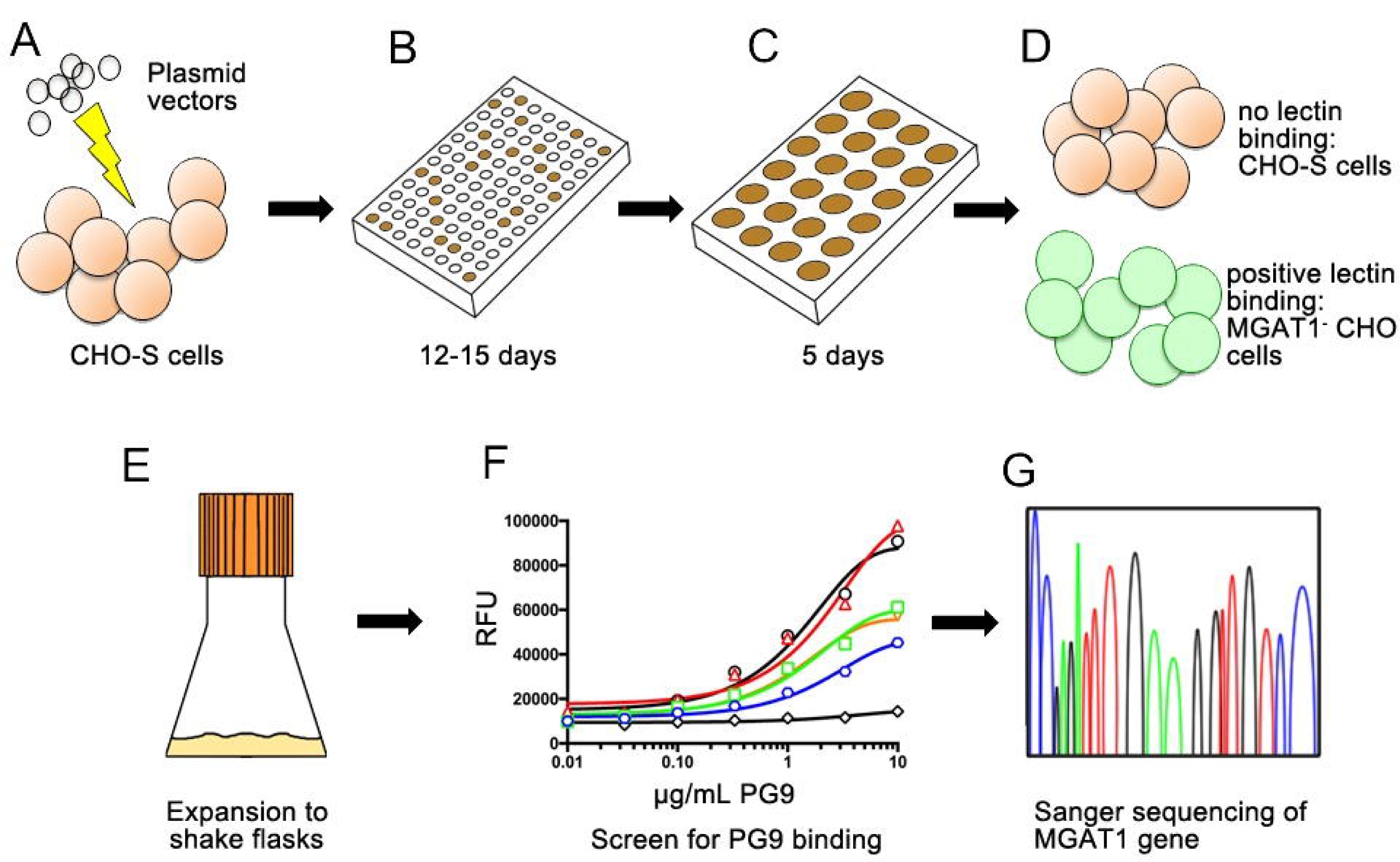
Flow chart of MGAT1 gene editing and cell line selection strategy. (A) A plasmid containing the Cas9 nuclease, tracrRNA, and a guide RNA (gRNA) sequence was electroporated into suspension adapted CHO-S cells. (B) Twenty-four hours following transfection, the cells were distributed into 96 well tissue culture plates at a density of 0.5 cells/well. (C) Between 12 and 15 days later, wells with 20% or greater confluency were transferred to 24 well plates. (D) After five days of growth in 24 well plates, a 0.2mL aliquot was removed from each well and cells were tested for the ability to bind fluorescein labeled Galanthus nivalis lectin (GNA). (E) GNA binding cells were then expanded to shake flasks and cell lines were transiently transfected with a gene encoding A244-rgp120. The cell culture supernatants were then collected after five days and tested for binding of gp120 to the prototypic glycan dependent, broadly neutralizing monoclonal antibody, PG9. (G) The gene encoding MGAT1 was sequenced from GNA binding cell lines with that exhibited robust growth and the ability to secrete PG9 binding gp120. The specific mutations induced by non-homologous end joining repair (NHEJR) were determined by Sanger sequencing.

### Lectin binding to detect MGAT1 gene inactivation

If the MGAT1 gene were inactivated, we expect glycoproteins to possess exclusively oligo-mannose forms of N-linked glycosylation, with a preponderance of Man_5_ isoforms on cell surface and secreted proteins. The lectin GNA recognizes glycans with terminal alpha-D mannose and is unable to bind to sialic acid containing complex glycans (48). Accordingly we used fluorescein conjugated GNA to determine whether CRISPR/Cas9 transfected cells possessed a phenotype characteristic of cells with an inactivated MGAT1 gene. GNA does not require Ca^2+^ or Mg^2+^ cofactors to bind, allowing the use of 10µM EDTA to ameliorate cell clumping during repeated centrifugation and wash steps. While MGAT1^-^ CHO cells and control HEK 293 GnTI^-^ cells bound to the GNA lectin, the wild type CHO-S cell line did not (Fig. 3). A total of 20 GNA binding cell lines from the original 166 candidates were selected on the basis of uniform GNA binding and the cultures were expanded for further analysis. Three days following initial GNA selection, the 20 cell line candidates were re-examined and six were rejected for lack of uniform lectin binding across the sample population, leaving 14 candidates.

**Fig 3.**
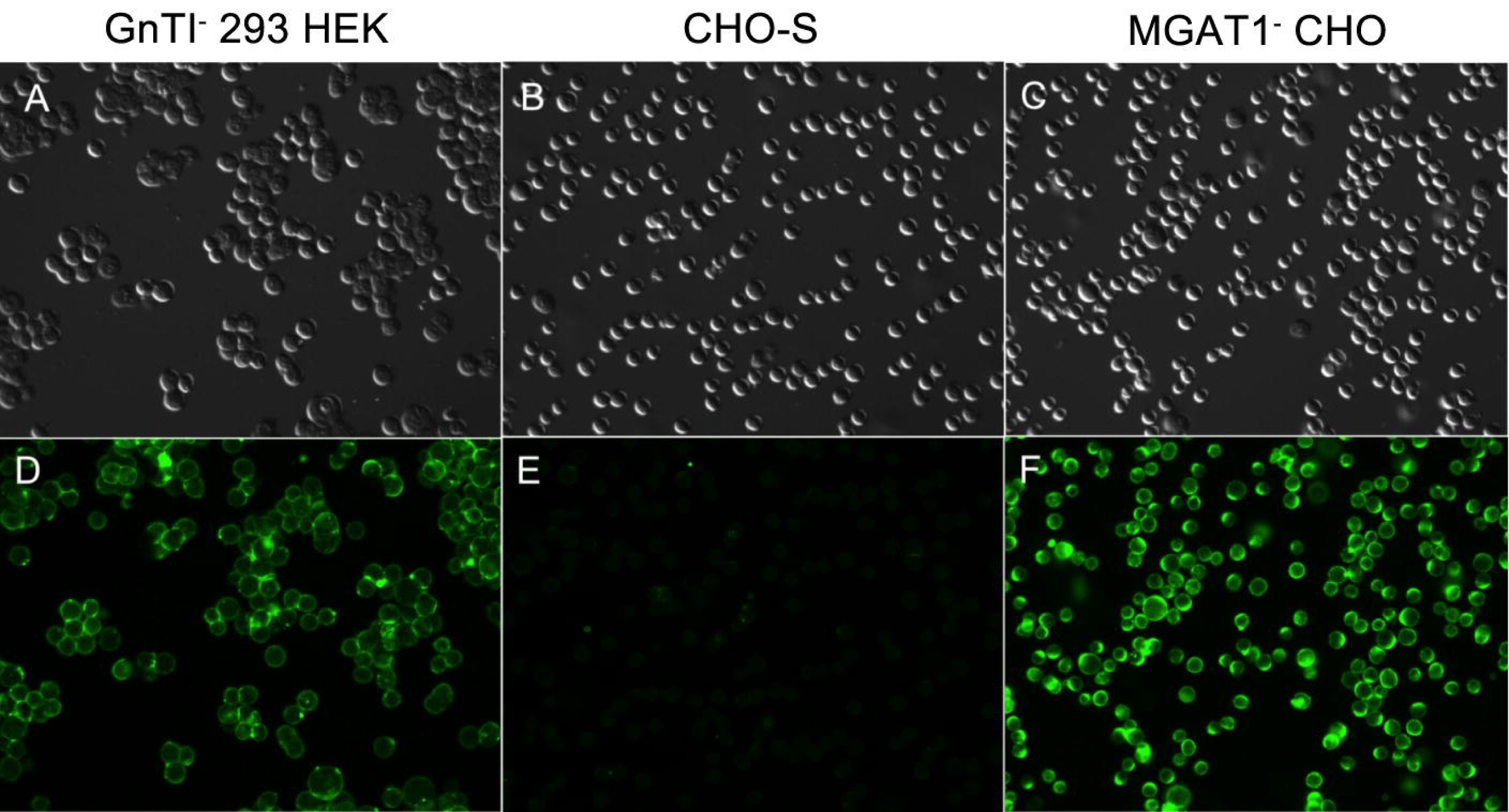
GNA lectin probe for cell surface oligomannose glycan expression. The GNA lectin binds glycan structures with terminal mannose and will not bind complex, sialic acid containing glycans. CHO-S cells were transfected with a plasmid designed to inactivate the MGAT1 gene by CRISPR/Cas9 gene editing (MGAT CHO). The cells were treated with fluorescein conjugated GNA lectin to screen for the incorporation of high mannose glycans in the cell membrane. HEK 293 GnTI^-^ cells that also lack the MGAT1 gene served as a positive control (panels A and D) while normal CHO-S cells that possess an intact MGAT1 gene served as a negative control (panels B and E). Cells were visualized under 20 x magnification on a Leica DM5500 B widefield microscope using differential interference contrast (DIC) (upper panels A, B, C) or under illumination with 495nm light (lower panels D, E, and F).

### Expression of gp120 in MGAT1^-^ CHO cell lines

Based on positive lectin binding criteria, the 14 candidate cell lines were grown in 125mL shake flasks (Fig. 2). Of those, the four fastest growing (3.4F10, 3.5D8, 3.5A2, and 3.4D9) were utilized for transient transfection with a gene encoding gp120 from the A244 strain of HIV-1 (A244-rgp120). Also transfected was the CHO-S parental cell line for comparison. This gp120 A244 had the sequence point mutations E332N and N334S, introducing a PNGS at N332. Five days post transfection the culture media was harvested and secreted gp120 proteins were purified by immunoaffinity chromatography. The purified products were assayed for protein yield, tested for identity using immunoblot analysis (data not shown) and for the ability to bind the glycan dependent bNAb, PG9 by fluorescent immune assay (FIA) (Fig. 4). Previous studies have shown that this bNAb requires Man_5_ at position N160 in the V1/V2 domain for binding (3). The results from this study confirmed that the MGAT1^-^ CHO cell lines could bind this antibody whereas gp120 produced in the parental CHO-S cell line was unable to bind PG9. From this analysis, a single MGAT1^-^ CHO cell line, 3.4F10, was selected for further characterization and analysis (Fig. 4).

**Fig 4.**
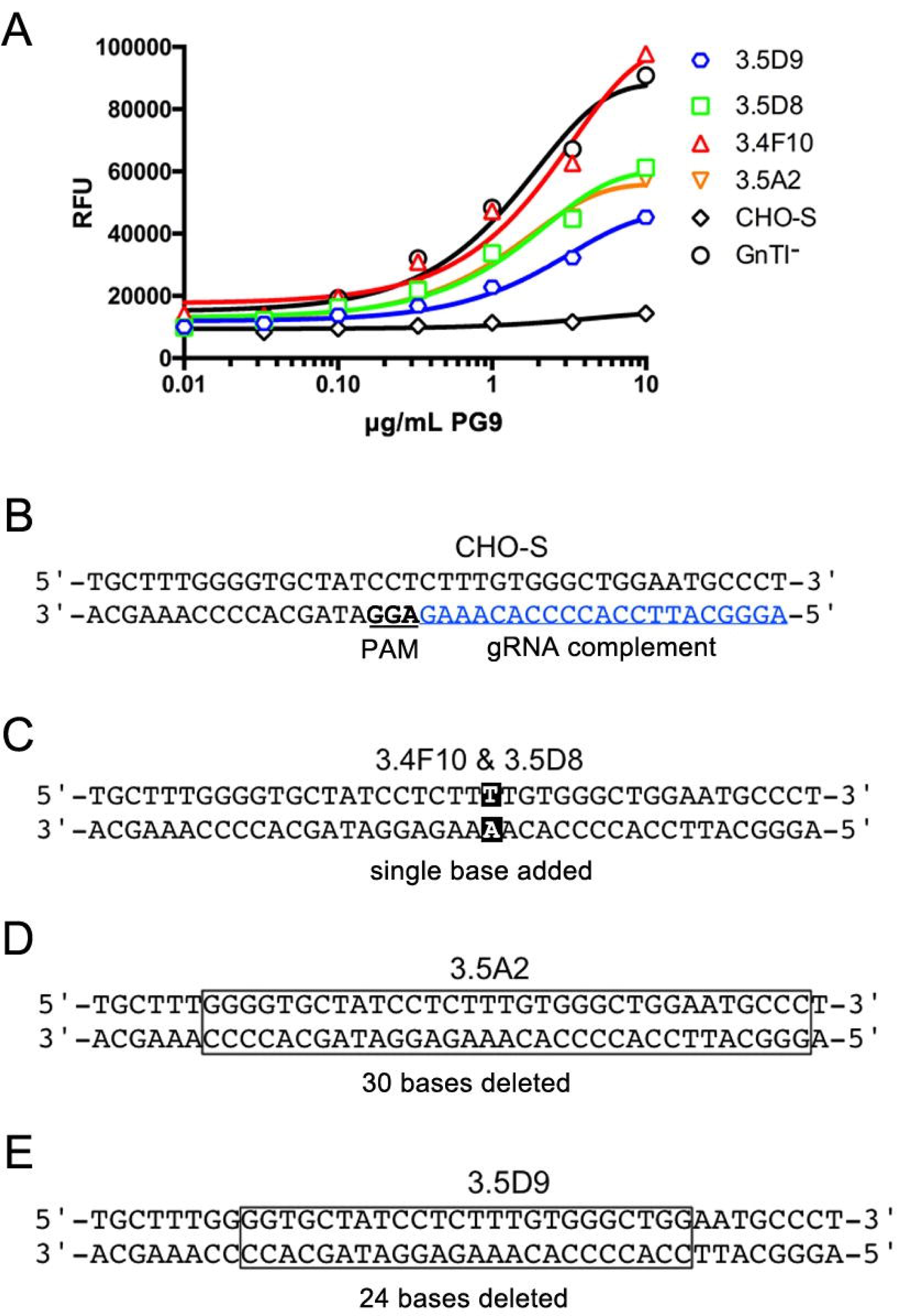
Screening and Sequence Analysis of MGAT1^-^ CHO cell line: Colonies selected after MGAT1 gene inactivation were transiently transfected with a gene encoding A244-rgp120. Cell culture supernatants were collected and tested for binding by the glycan dependent bN-mAb PG9. Based on PG9 binding studies the MGAT genes from selected cell lines were amplified by PCR and sequenced. (A) PG9 binding to gp120 in cell culture supernatants of transiently transfected MGAT1^-^ CHO lines 3.5D9, 3.5D8, 3.4F10, 3.5A2, and by supernatants from gp120 transfected CHO-S and GnTI^-^ 293 HEK cells. (B) Diagram of the unaltered CHO-S MGAT1 gene target section with guide RNA (gRNA) complement sequence shown in blue and the protospacer adjacent motif (PAM) underlined in bold type. (C) Sequences of the MGAT1 gene for 3.4F10 and 3.5D8 cell lines both had the same single base insertion, shown in black box. (D) The sequence from the cell line 3.5A2 with bases deleted shown in box. (E) The bases deleted 3.5D9 cell line sequences are indicated by the box.

To be a viable substrate for biopharmaceutical production, the growth and protein yield of the knockout line had to be comparable to the parental line. In transient transfection experiments the calculated recovery of purified protein was 35.4mg/L for the 3.4F10 MGAT1^-^ CHO line and 32.2mg/L for the parental CHO-S line. Production of the same protein in HEK 293 GnTI^-^ cells by transient transfection yielded 1.9mg/L. We measured that the cell doubling time for the 3.4F10 MGAT1^-^ CHO cell line in BalanCD CHO Growth A media was 20.7 hours in a 1L shaker flask during logarithmic growth phase, reaching a density of 1.9×10^7^ cells/mL. This was similar to the parental CHO-S population doubling time of 19.0 hours that achieved cell densities of 1.6×10^7^ cells/mL. By comparison, the GnTI^-^ HEK 293 cell line had a logarithmic cell doubling time of 23.3 hrs. and achieved a maximal cell density of 4.20 x10^6^ cells/mL when grown in Freestyle 293 media.

### Identification of CRISPR/Cas9 induced genetic alteration

To confirm that MGAT1 gene had been inactivated, we sequenced the gene from the 3.4F10 line and the next three best candidates. An extra thymidine had been inserted at the Cas9 cleavage site of the 3.4F10 line MGAT1 gene, introducing a frame shift mutation. This mutation resulted in 23 altered codons and the insertion of a premature stop codon. The 3.5D8 line contained the same mutation, while 3.5D9 and 3.5A2 both had in frame deletions of 24 and 30 nucleotides respectively. The deleted codons of 3.5D9 and 3.5A2 corresponded to the transmembrane domain of the GnTI protein leaving the active extracellular domain intact. The diminished binding of gp120s produced in the 3.5D9 and 3.5 A2 clones to PG9 suggest that partial MGAT1 activity remains in these two clones compared to the 3.4F10 clone that, like gp120 produced in GnTI^-^ cells, exhibits improved binding to PG9 (Figure 4). Given that the single base insertion in the 3.4F10 MGAT1 gene resulted in a frame shift 51 nucleotides into a 1276bp long gene, it is highly unlikely that the function of this gene could be restored by random mutation.

### Characterization of MGAT1^-^ CHO gp120 glycosylation

Two additional methods (endoglycosidase digestion and mass spectrometry analysis using MALDI-TOF-MS) were used to further characterize the N-linked glycosylation incorporated in A244-rgp120 produced by the 3.4F10 MGAT1^-^ CHO cell line. Immunoaffinity purified, monomeric A244 gp120 produced by the CHO-S, HEK 293 GnTI^-^, and MGAT1^-^ CHO cell lines were digested overnight by endoglycosidases PNGase F and Endo H, then analyzed by SDS-PAGE and stained with Coomassie blue dye (Fig. 5). Endo H cleaves N-linked high-mannose glycan structures, and not complex, sialic acid containing glycans. When the protein produced in the HEK 293 GnTI^-^ and MGAT1^-^ CHO cell lines was compared to the proteins produced in CHO-S cell lines, we noted a reduction in mass of approximately 20kD. This is in keeping with the smaller mass of the Man_5_ glycoform compared to that of the hybrid and complex glycans found on CHO-S produced material. Following Endo H digestion, the protein produced in the CHO-S cell line was largely unaltered, indicating that it possessed the normal complex, sialic acid containing glycans. In contrast, the proteins produced in the MGAT1^-^ CHO and HEK 293 GnTI^-^ cells were reduced to ∼60kD in size. This result was consistent with the observation that approximately half the mass of a given gp120 molecule can be attributed to N-linked glycosylation (47, 49, 50). The complete sensitivity of the proteins produced in the MGAT1^-^ CHO and HEK 293 GnTI^-^ cells to Endo H digestion suggests that the glycosylation of these cell lines is exclusively high-mannose. When digested with PNGase F, all samples dropped to the same size, confirming that undigested gp120 size variances were due to glycosylation differences.

**Fig 5.**
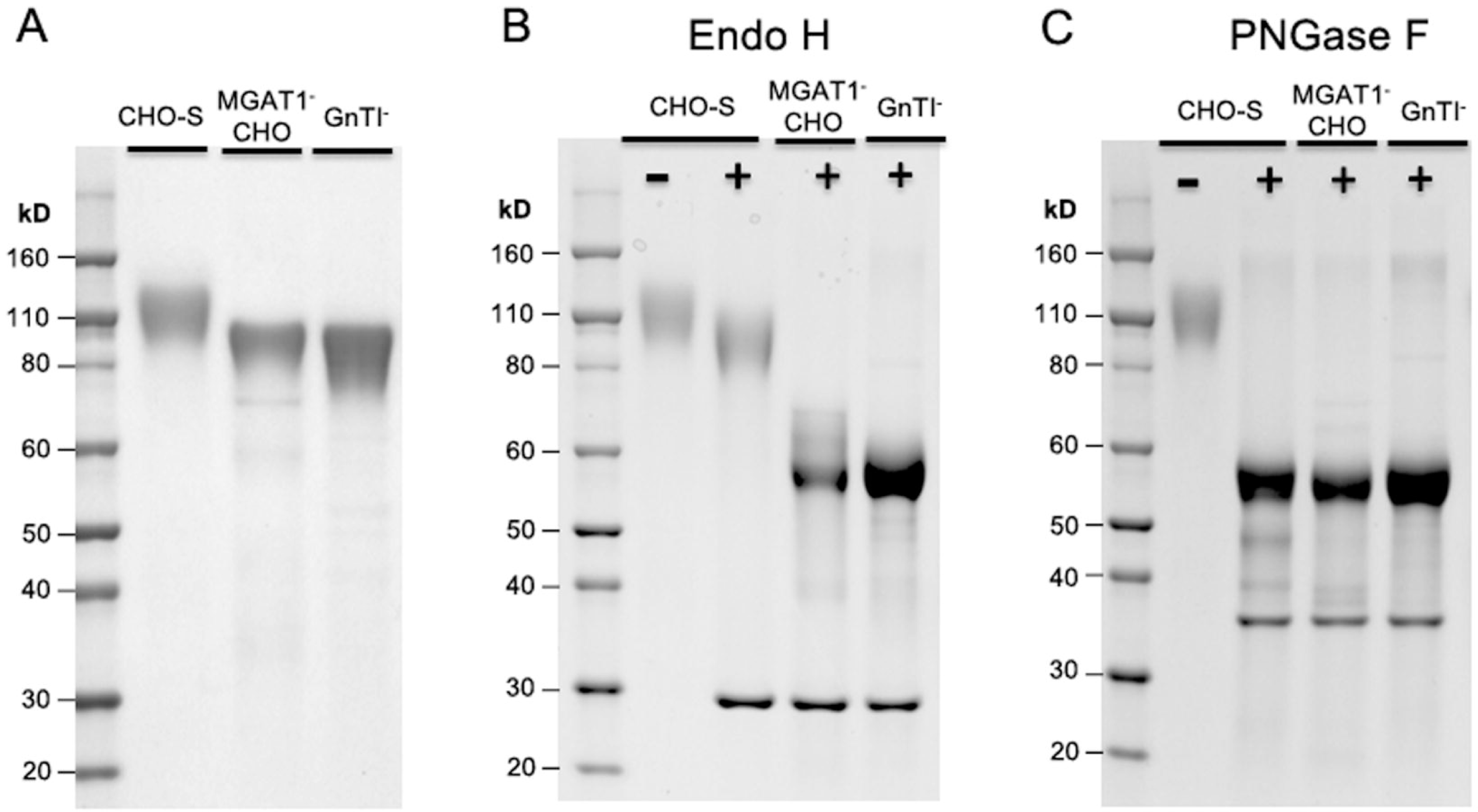
Endoglycosidase analysis of gp120 produced in MGAT1^-^ CHO cell line. Purified A244 rgp120 recovered from transiently transfected CHO-S, MGAT1^-^ CHO, or HEK 293 GnTI^-^ cell lines was analyzed by SDS-PAGE following endoglycosidase treatment. Purified gp120s were reduced and denatured then treated with either endoglycosidase H (Endo H) or Peptide:N-Glycosidase F (PNGase F). The digests were then analyzed on 4–12% tris-glycine SDS PAGE gels and stained with Coomasie blue dye. Panel A, mock digests of gp120s produced in CHO-S, MGAT1^-^ CHO, and HEK 293 GnTI^-^ cells. Panel B, the same proteins in panel A, digested with endoglycosidase H (Endo H). Panel C, the same proteins in panel A, digested with PNGase F. The mobility of molecular weight markers is shown for each gel. The endoglycosidases proteins are visible as bands at 29kD (Endo H, panel B) and 36kD (PNGase F, panel C).

Additional studies were carried out to characterize the specific glycans incorporated in the A244-rgp120 produced in the MGAT1^-^ CHO and the CHO-S cell lines. Using MALDI-TOF-MS (Fig. 6), we found that 56.4% of the N-linked glycans present on the MGAT1^-^ CHO produced gp120 were Man_5_, 19.2% were Man_9_, 11% were Man_8_ and the remainder were Man_6_ and Man_7_. No complex sialic acid containing glycans were detected (Table 1). The degree of fucosylation was also significantly lowered; fucosylation was only on Man_5_ glycoforms at the core GlcNAc and represented 3.16% of the total glycans present.

**Fig 6.**
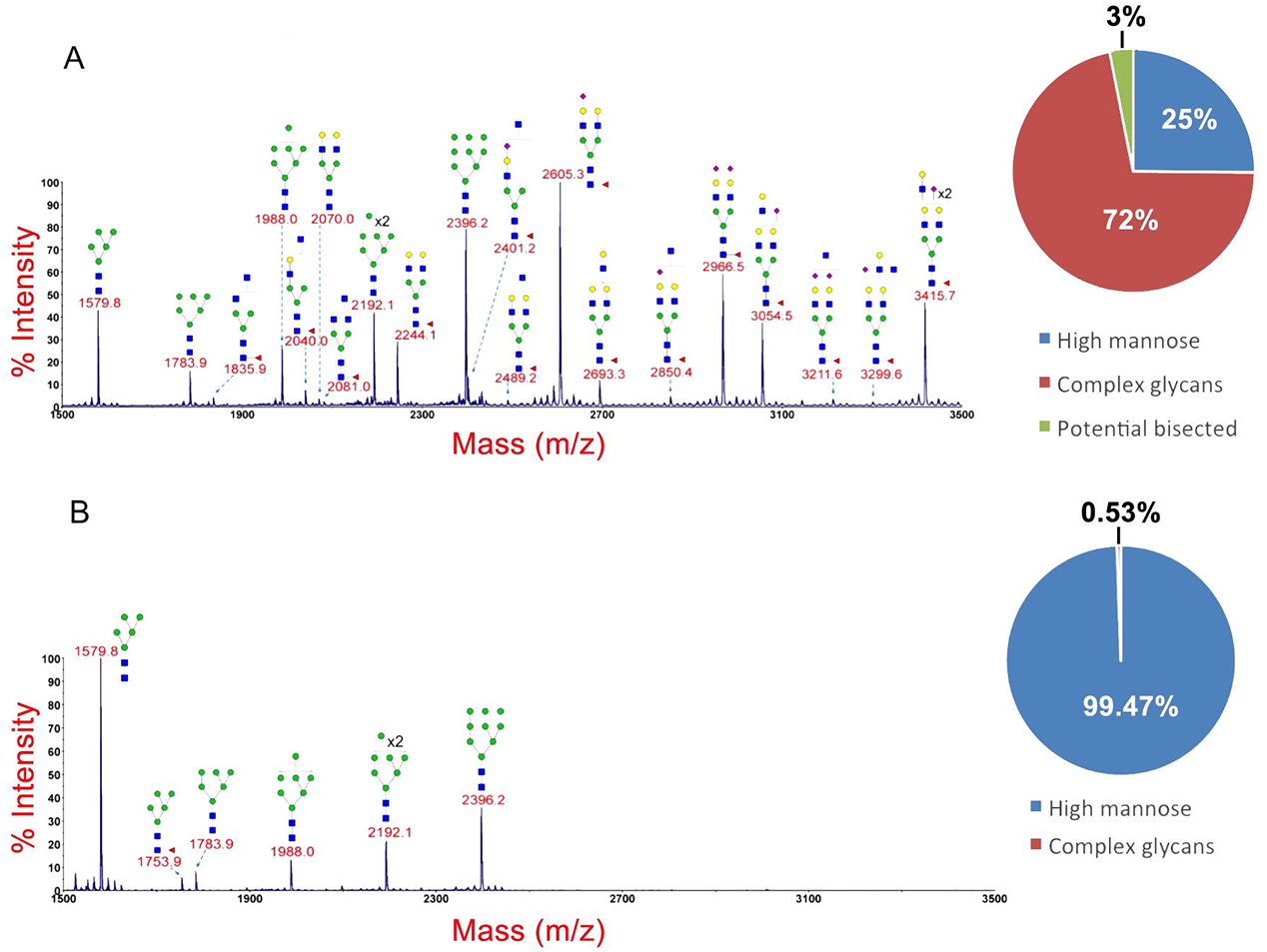
MALDI-TOF analysis of glycans present on gp120 produced by CHO-S and MGAT1^-^ CHO cell lines. The carbohydrates on purified A244 gp120s produced by CHO-S (A) and MGAT1^-^ CHO (B) cells were released by PNGase F digestion and examined by MALDI-TOF MS as described in Materials and Methods. Pie charts indicate the percentage of high-mannose (blue), complex (red), and potential bisected (green) N-linked glycans. This analysis was performed by the Complex Carbohydrate Research Center at the University of Georgia.

**Table 1.** Percentage of different glycan species on gp120 produced by CHO-S and MGAT1^-^ CHO cells.

| Glycan Species | Cell Line |  |
| --- | --- | --- |
|  | CHO-S | MGAT1<sup>-</sup> CHO |
| Man<sub>5</sub> | 5.2% | 56.4% |
| Man<sub>6</sub> | 1.9% | 4.5% |
| Man<sub>7</sub> | 3.3% | 7.1% |
| Man<sub>8</sub> | 5.1% | 11.4% |
| Man<sub>9</sub> | 9.6% | 19.2% |
| Complex or Hybrid | 75% | 0.53% |

Percentages calculated from MALDI-TOF peak intensity.

When the A244-rgp120 produced in CHO-S cells was examined, approximately 75% of the glycans were complex or hybrid glycans and 25% represented the early intermediates ranging from Man_5_ to Man_9_. No high mannose species were detected with core GlcNAc fucose attached, but nearly all hybrid and complex glycans were fucosylated.

### Binding multiple bN-mAbs to gp120 expressed in the MGAT1^-^CHO cells

We next compared A244-rgp120 expressed in the MGAT1^-^ CHO cells with A244-rgp120 produced in normal CHO-S cells for the ability to bind bN-mAbs in a FIA. A panel of prototypic bN-mAbs that recognize distinct sites of virus vulnerability in monomeric and trimeric HIV envelope proteins were utilized (Fig. 7). We noted a significant improvement in the binding of PG9, CH01 and CH03 to the proteins expressed in MGAT1^-^ CHO cells and HEK 293 GnTI^-^ cells compared to the CHO-S cells. These bN-mAbs are known to bind to epitopes in the V1/V2 domain that require Man_5_ at the N160 glycosylation site (3). Similarly we noted a significant improvement in the binding of the PGT126 and PGT128 bN-mAbs that require oligo-mannose glycans at the N301 and N332 glycosylation sites in the stem of the V3 domain (51). Mixed results were seen for the PGT121 family of bN-mAbs where the binding to gp120 produced in both the MGAT1^-^ CHO and HEK 293 GnTI^-^ cells lines was lower than binding of these antibodies to gp120 produced in the CHO-S cell line. In contrast, binding to the 10–1074 bN-mAb, also in the PGT121 family, was unaffected by the cellular substrate used for production. These results demonstrate that changing the glycosylation, while leaving the amino acid sequence intact can significantly improve the antigenic structure of A244-rgp120 with respect to the binding of several bNAbs to glycan dependent epitopes. The effect of differences in glycosylation was also examined on the binding of the VRC01 bN-mAb, known to recognize a glycan independent epitope adjacent to the CD4 binding site (52). While this antibody bound to all of the envelope proteins tested, a small, but consistent improvement in binding was observed to the proteins produced in the MGAT1^-^ CHO and HEK 293 GnTI^-^ cells compared to the protein produced in the CHO-S cells. These studies suggest that the sialic acid containing hybrid and complex carbohydrates incorporated in normal cell lines in some way interfere with the VRC01 binding site on monomeric gp120. This same effect may be observed with further non-glycan dependent antibodies by decreasing the glycan interference.

**Fig 7.**
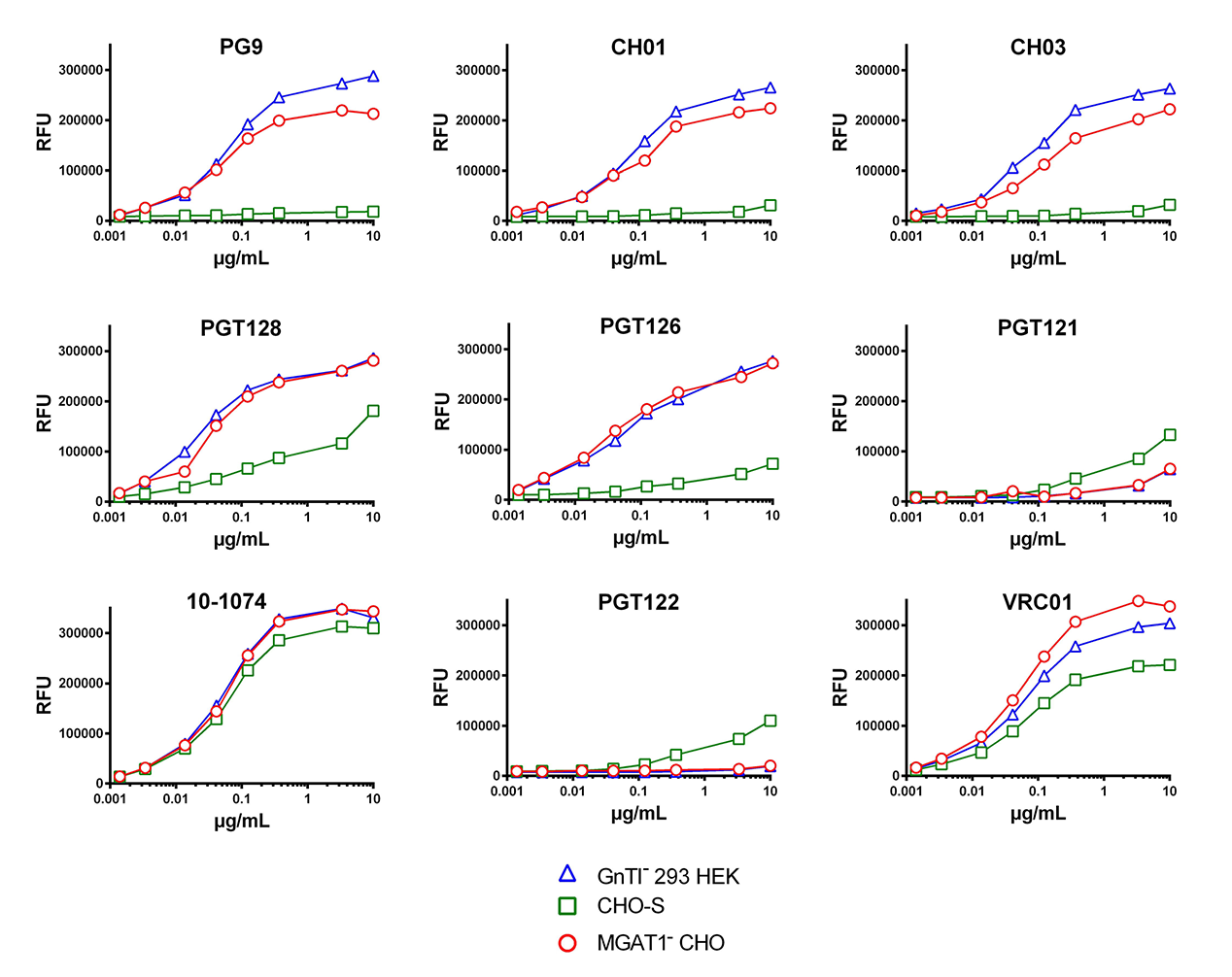
Comparison of bN-mAb binding to A244-rgp120 produced in MGAT1^-^ CHO cells, CHO-S cells and HEK 293 GnTI^-^ cells. The binding of a panel of broadly neutralizing monoclonal antibodies to purified A244-rgp120 produced in MGAT1^-^ CHO cells, CHO-S cells, and HEK 293 GnTI^-^ cells was measured in a Fluorescence Immunoassay (FIA). Briefly, purified proteins were captured onto wells of black 96 well microtiter plates coated with a mouse monoclonal antibody against the N-terminal gD tag present in all three proteins. Plates were then incubated with serial dilutions of bN-mAbs targeting: glycan-epitopes within the V1V2 domain (PG9, CHO1 and CHO3), the glycan-epitopes within the V3 domain (PGT128, PGT126, PGT121, 10–1074 and PGT122), or the CD4 binding site (VRC01). Plates were incubated with a 1:3,000 dilution of AlexaFluor 488 conjugated goat-anti-human polyclonal antibody and binding is reported as Relative Fluorescence Units (RFU). FIA details are provided in Materials and Methods.

### MVM infectivity

Minute virus of mice (MVM) is a small inactivation resistant virus that is ubiquitous in the environment and a major cause of bioreactor culture failure in biopharmaceutical manufacturing (53). As sialic acid is a major receptor for MVM infectivity, the MGAT1^-^ CHO cell line we created might have the additional manufacturing benefit of being resistant to MVM infection (54). To investigate this possibility, the MGAT1^-^ CHO cell line was tested for infectivity resistance to two strains of MVM using a qPCR assay and compared to wild type CHO-S MVM sensitivity. While the MGAT1^-^ CHO cell line was similarly sensitive to the MVMp strain as wild type CHO-S cells, it was resistant to MVMc infection (Table 2). The receptor protein for MVM have not yet been identified, but it has been demonstrated that MVMp binds to sialic acid residues from both N and O linked glycosylation (55–57). Knocking out MGAT1 does not alter the O-linked glycosylation pathway, perhaps explaining why the line remains sensitive to MVMp. MVMc is a more recently identified strain(58) with little information available on its binding to CHO or murine cells. Anything beyond noting the apparent dependence on complex N-linked glycosylation would be speculative at this point.

**Table 2:**
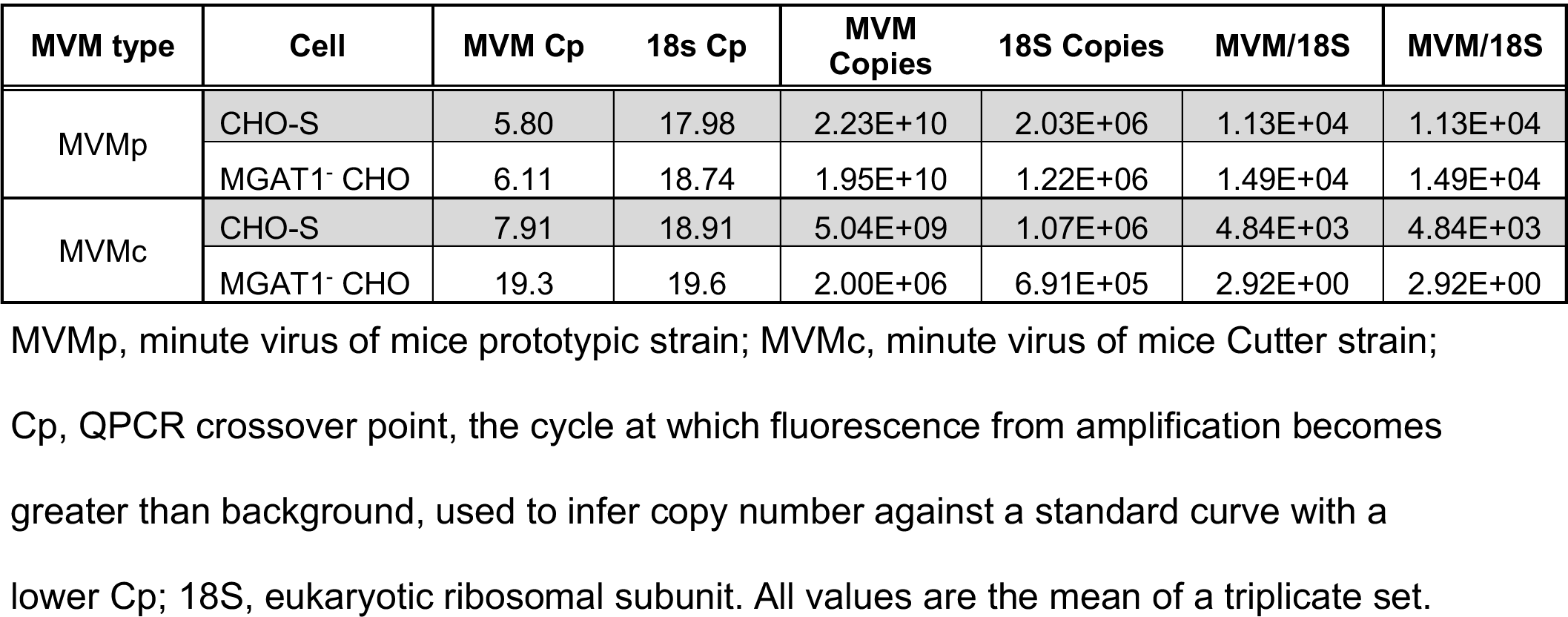
MVM infectivity assay.

This infectivity assay was performed by IDEXX BioResearch (Columbia, MO).

## Adventitious agent testing

The cell line was tested for the presence of mycoplasma, cross-species contamination, and viral contaminants by IDEXX BioResearch (Columbia, MO). No adventitious agents were detected. The full list and procedure are described in the supplemental materials.

## Discussion

A major goal in HIV vaccine research is to develop immunogens that elicit bNAbs. The discovery that multiple bN-mAbs to HIV recognize glycan dependent epitopes has altered our thinking of how best to produce this vaccine. Instead of using standard CHO cell production cell lines, that incorporate complex and hybrid glycosylation, a cell line that limits glycosylation to high-mannose forms may be useful for gp120 immunogens. While we have long been able to produce properly folded Env protein monomers (gp120 and gp140) as indicated by the ability to bind CD4 with high affinity (59–63), until now we have not been able to replicate the glycan structures required for the binding of multiple families of bN-mAbs in expression systems suitable for large scale manufacturing.

The glycans that decorate the surface of native, virion-associated, HIV Env protein are typically enriched for high-mannose variants, normally found on early intermediate proteins within the ER and early Golgi (18, 19). This unusual restriction in glycan maturation is thought to be a consequence of steric hindrance occurring during the formation of trimeric virus spike structures (64). Additionally, the high density of PNGS that likely evolved as a glycan shield to prevent immune recognition of virus sequences (38, 47, 65) appears to limit glycosidase and glycosyltransferase modifications of Env glycans in the late ER and early Golgi Apparatus (GA) (18, 64). While expression of monomeric gp120 results in incorporation of complex glycosylation, trimeric spike formation results in incomplete glycosylation and enrichment of virions with high-mannose glycans (18, 19). These differences in glycosylation might explain the inability of previous HIV vaccines to elicit bNAbs to glycan dependent epitopes in humans. However they don’t explain the inability of previous vaccines to elicit bNAbs to epitopes such as VRC01 that were present in most gp120 vaccines expressed in normal CHO cells. Earlier vaccines such as the AIDSVAX B/E used in the RV144 trial largely possessed complex sialic acid containing glycans and lacked the high-mannose glycans required for a variety of bN-mAbs including PG9, CH01, CH03, PGT128, and 10–1074 (16). Although the level of protection achieved in the 16,000-person RV144 trial was statistically significant (31.2%, P=0.04) this level was not sufficient for product registration or clinical deployment. In this regard, the addition of one or more epitopes recognized by bN-mAbs, such as the PG9 and PGT128 epitopes described in this report, might improve the antigenic structure and immunogenicity of or recombinant gp120 such that a level of protection of 50% or more, required for product approval is achieved. Currently RV144 follow-up studies are in progress that make use of sialic acid containing gp120 vaccine antigens, produced in normal CHO cells, like those used in the original RV144 trial (66, 67). These new trials are trying to improve the level of vaccine efficacy by prolonging the immunization schedule, altering the adjuvant formulation, and replacing the canarypox vector co-administered with gp120 with stronger, more virulent virus vectors.

Few methods currently exist to produce recombinant proteins incorporating the Man_5_ and Man_9_ glycans that are present in the HIV gp120 envelope protein. Expression of gp120 in yeast results in the incorporation of long chain high-mannose glycans (68), and insect expression systems produce a preponderance of paucimannose forms (Man_3–4_) (69). Glycosidase inhibitors (e.g. kifunensine and swainsonine, see Fig. 1) are effective and useful for producing analytical quantities of proteins with Man_5_ and Man_9_ intermediates, but are highly toxic and prohibitively expensive for large-scale biopharmaceutical production (70, 71). Additionally, there exists evidence to suggest a broad mannosidase inhibitor like kifunensine may negatively impact protein folding through interference of the Calnexin/Calreticulin pathway (72–75). Glycosaminyl-transferase knockout cell lines from 293 HEK and CHO cells, referred to as HEK 293 GnTI^-^ and CHO Lec1, respectively, have previously been described. They were generated through random ethyl methanesulfonate (EMS) mutagenesis, zinc finger methods, or screened for modified glycosylation by resistance to cytotoxic lectin binding (76–78). These lines lack a functional MGAT1 gene, responsible for the protein GnTI. Knocking out the MGAT1 gene prevents processing of glycans beyond the Man_5_GlcNAc_2_ stage, resulting in exclusively high-mannose glycoprotein production (28, 29). Such cell lines do not generally grow as robustly as their parental counterparts and raise potential regulatory issues with potential uncharacterized genetic alterations. In light of this, there is an unmet need for a cell line suitable for the scalable production of HIV vaccine immunogens. We have addressed this problem by creating the novel MGAT1^-^ CHO cell line described above. Our data suggests that this cell line possesses several essential characteristics required for current Good Manufacturing Practices (cGMP) such as robust growth in well defined serum free medium, the ability to grow to high cell densities in suspension culture, a well defined mutation of the MGAT1 gene, and freedom from contamination by adventitious agents. However, the ultimate utility of this cell line will require characterization of stable cell lines with transfected envelope proteins where the genetic stability of the transgene as well as the quality and yield of the final product is determined.

Recombinant envelope proteins produced in the MGAT1^-^ CHO cell line, such as the A244-rgp120 described in this report, can be used to test the hypothesis that previous HIV vaccines such as the AIDSVAX B/E vaccine used in the RV144 trial (12–14) were ineffective because they lacked the glycan dependent epitopes required to stimulate the formation of bN-mAbs. While the CHO MGAT1^-^ cell line provides a practical way to produce envelope proteins with several glycan dependent epitopes recognized by bNAbs not present on gp120s produced in normal CHO cell lines, we do not yet know whether these epitopes will be immunogenic. Previous gp120 vaccine trials such as RV144 failed to detect glycan independent VRC01-like antibodies even though the VRC01 epitope was present on at least two different gp120s used for immunization. Thus, some epitopes recognized by bNAbs are poorly immunogenic and additional immunogenicity and formulation studies will likely be required to optimize virus neutralizing antibody responses to the glycan epitopes of the type described in this paper. Although the gp120 expression data presented here was derived exclusively from small-scale transient transfection experiments, we anticipate that the cell line described in this report will be useful for the development of stable MGAT1^-^ CHO cell lines producing vaccines based on a variety of new concepts. These include guided immunization to stimulate germline genes encoding bNAbs (79–81), Env proteins designed with features that enhance antigen processing and presentation (82) as well as glycopeptide scaffolds that enhance the immunogenicity of epitopes recognized by bN-mAbs while eliminating non-protective immunodominant epitopes (21).

## Materials and methods

### Cell culture

Suspension adapted CHO-S cells were obtained from Thermo Fisher (Thermo Fisher, Life Technologies, Carlsbad, CA). HEK 293 GnTI^-^ suspension adapted cells were obtained from ATCC (ATCC, Manassas, VA). Stocks of suspension adapted CHO-S and HEK 293 GnTI^-^ cells were maintained in shake flasks (Corning, Corning, NY) using a Kuhner ISF1-X shaker incubator (Kuhner, Birsfelden, Switzerland). For cell propagation, shake flask cultures were maintained at 37°C, 8% CO_2_, and 125 rpm. Static cultures were maintained in 96 or 24 well cell culture dishes and grown in a Sanyo incubator (Sanyo, Moriguchi, Osaka, Japan) at 37°C and 8% CO_2_.

CHO-S cells were maintained in CD-CHO medium supplemented with 0.1% pluronic acid, 8mM GlutaMax and 1X Hypoxanthine/Thymidine (Thermo Fisher, Life Technologies, Carlsbad, CA). For cell growth studies, CHO cells were grown in BalanCD CHO Growth A Medium (Irving Scientific, Santa Ana, CA). HEK 293 GnTI^-^ cells were maintained in Freestyle 293 cell culture media (Life Technologies, Carlsbad, CA). During transient CHO cell protein production the cells were maintained in OptiCHO medium supplemented with 0.1% pluronic acid, 2mM GlutaMax and 1X H/T (Thermo Fisher, Life Technologies, Carlsbad, CA). For protein production experiments the growth medium was supplemented with CHO Growth A (Molecular Devices, Sunnyvale, CA), 0.5% Yeastolate (BD, Franklin Lakes, NJ), 2.5% CHO-CD Efficient Feed A; and 0.25mM GlutaMax, 2 g/L Glucose (Sigma-Aldrich St. Louis, MO). Cell counts were performed using a TC20^TM^ automated cell counter (BioRad, Hercules, CA) with viability determined by trypan blue (Thermo Fisher, Life Technologies, Carlsbad, CA) exclusion.

Cell-doubling time in hours was calculated using the formula: ((T_2_-T_1_) × log_2_)/(log(D_2_)log(D_1_)), where T = time at count and D = density at count. Cell count numbers used for doubling time calculation were from the logarithmic phase of growth.

### Gene sequencing

The sequence of the MGAT1 CHO gene was confirmed using primers based on the predicted mRNA transcript (XM_007644560.1 (83)). Genomic DNA was extracted using the AllPrep kit (Qiagen, Germantown, MD). The MGAT1 gene was PCR amplified using the primers F_CAGGCAAGCCAAAGGCAGCCTTG and R_CTCAGGGACTGCAGGCCTGTCTC (Eurofins Genomics, Louisville, KY) with Taq and dNTPs supplied by New England BioLabs (Ipswich, MA). The PCR product was gel purified using a Zymoclean kit (Zymo Research, Irvine, CA), then sequenced by Sanger method at the (UC Berkeley, Berkeley, CA). MGAT1 knockouts were sequenced in the same manner.

### CRISPR/Cas9 target design and plasmid preparation

We utilized a CRISPR/Cas9 nuclease vector with an OFP reporter (GeneArt, Thermo Fischer Scientific, Waltham, MA). Three target sequences to knock out the CHO-S MGAT1 gene were designed using an online CRISPR RNA Configurator tool (GE Dharmacon, Lafayette, CO). Target 1: CCCTGGAACTTGCGGTGGTC.Target 2: GGGCATTCCAGCCCACAAAG. Target 3: GGCGGAACACCTCACGGGTG. Each sequence was run in NCBI’s BLAST tool for homologies with off-target sites in the CHO genome. Single stranded DNA oligonucleotides and their complement strands were synthesized (Eurofins Genomics, Louisville, KY) with extra bases on the 3’ ends for ligation into GeneArt CRISPR nuclease vector (Thermo Fisher, GeneArt, Waltham, MA).

The strands were ligated and annealed into a GeneArt CRISPR vector using the protocol and reagents supplied with the kit. One Shot^®^ TOP10 Chemically Competent *E. coli* were transformed and plated following the Invitrogen protocol (Thermo Fisher, Invitrogen, Carlsbad, CA). These were incubated in 5mL LB broth at 37°C in a shaking incubator at 225rpm overnight. Minipreps were performed according to manufactures instructions (Qiagen, Germantown, MD) and sent to UC Berkeley DNA Sequencing Facility (Berkeley, CA) with the U6 primers included in the GeneArt^®^ CRISPR kit to confirm successful integration of guide sequences. A single 500mL Maxiprep was performed for each of the three target sequences using PureLink^tm^ Maxiprep kit (Thermo Fisher, Invitrogen, Carlsbad, CA).

### Electroporation

Electroporation of CHO cells was performed using a MaxCyte STX scalable transfection system (MaxCyte Inc., Gaithersburg, MD) according to the manufacturer’s instructions. Briefly, CHO-S cells were maintained at >95% viability prior to transfection. Cells were pelleted at 250g for 10 minutes, and then re-suspended in MaxCyte EP buffer (MaxCyte Inc., Gaithersburg, MD) at a density of 2×10^8^ cells/mL. Transfections were carried out in the OC-400 processing assembly (MaxCyte Inc., Gaithersburg, MD) with a total volume of 400µL and 8×10^7^ total cells. CRISPR/Cas9 exonuclease with guide sequence plasmid DNA suspended in endotoxin-free water was added to the cells in EP buffer for a final concentration of 300μg of DNA/mL. The processing assemblies were then transferred to the MaxCyte STX electroporation device and CHO protocol was selected using the MaxCyte STX software. Following electroporation, the cells in electroporation buffer were removed from the processing assembly and placed in 125mL Erlenmeyer cell culture shake flasks (Corning, Corning, NY). The flasks were placed into 37°C incubators with no agitation for 40 minutes. Following the rest period pre-warmed OPTI-CHO media was added to the flasks for a final cell density of 4×10^6^ cells/mL. Flasks were then moved to a Kuhner shaker and agitated at 125rpm.

### Plating, expansion, and culture of CRISPR transfected CHO-S cells

Twenty-four hours post transfection a 100μL aliquot was taken from each of the transfected pools to assay for cell viability and orange fluorescent protein (OFP) expression using a light microscope (Zeiss Axioskop 2, Zeiss, Jena, Germany). Ninety-six well flat bottom cell culture plates (Corning, Corning, NY) were filled with 50μL of conditioned CD-CHO media. Each of the three transfected pools were serially diluted with warmed media to 10 cells/mL and added to five plates per pool in 50μL volumes. Final calculated cell density was 0.5 cells/well in 100μL of media. Once any single-colony well reached ≈20% confluency, the contents were transferred to a 24 well cell culture plate (Corning, Corning, NY) with 500uL of fresh media. When confluency reached 50%, a 200μL aliquot was removed for testing via a GNA lectin-binding assay. Following positive lectin binding, cells were moved to a 6 well cell culture plate (Corning, Corning, NY) with 2mL of media per well. After 5 days of growth in 6 well plates the GNA assay was repeated. Colonies exhibiting positive lectin binding were moved to 125mL shake flasks with an initial 6mL of media. Daily counts were taken and cell cultures expanded to maintain 0.3×10^6^ – 1.0×10^6^ cells/mL density.

### Lectin binding assay

Fluorescein labeled *Galanthus nivalis* lectin (GNA), from the snowdrop pea (Vector Laboratories, Burlingame, CA), was used to detect the cell surface expression of Man_5_ glycoforms. Cell aliquots (200μL) from 24 well plates were pelleted at 3000 rpm for three minutes. The supernatant was discarded and the cell pellet washed three times with 500μL of ice-cold 10μM EDTA in (Boston BioProducts, Ashland, MA) phosphate buffered saline (PBS) (Thermo Fisher, Gibco, Carlsbad, CA). The cell pellet was then re-suspended in 200μL ice cold 10μM EDTA with PBS with 5μg/mL of GNA-fluorescein. Samples were shielded from light and incubated on ice with GNA for 30 minutes. Following incubation, samples were washed three times and re-suspended to a volume of 50μl in 10μM EDTA PBS. Samples were then examined under a light microscope (Zeiss Axioskop 2, Zeiss, Jena, Germany) with 495nm wavelength excitation. Wild type CHO-S cells were used as a negative control and HEK 293 GnTI^-^ were used as a positive control. Representative images were taken on a Leica DM5500 B Widefield Microscope (Leica Microsystems, Buffalo Grove, IL) at the UC Santa Cruz microscopy center.

### Experimental protein production

An expression plasmid containing the gene encoding gp120 from the A244 strain of HIV (Genbank accession number: MG189369) was selected for transient transfection experiments. The protein encoded by this gene was identical to that used to produce the AIDSVAX B/E vaccine used in the RV144 trials (13, 14) with the exception that the N-linked glycosylation site at N334 was moved to N332. For analytical scale experiments, a total of 4×10^5^ cells from each candidate MGAT1^-^ CHO line were placed in 450μl of media in a 24 well cell culture plate. Fugene, 1.7μL, (Promega, Madison, WI) was pre-incubated at room temperature for 30 minutes with 550ng of DNA in a total volume of 50μL of media. Then 50μL of Fugene/DNA mixture was added to each well for a final transfected volume of 500μL. Aliquots of supernatant were removed for assay 72 hours post transfection.

For preparative scale transient transfection experiments, CHO cells were electroporated following the above MaxCyte method. Twenty-four hours post electroporation, the culture was supplemented with 1mM sodium butyrate (Thermo Fisher, Life Technologies, Carlsbad, CA) and temperature lowered to 32°C. The cultures were fed daily the equivalent of 3.5% of the original volume with Molecular Devices CHO A Feed (Molecular Devices, Sunnyvale, CA), 0.5% Yeastolate (BD, Franklin Lakes, NJ), 2.5% CHO-CD Efficient Feed A; and 0.25mM GlutaMax, 2 g/L Glucose (Sigma-Aldrich St. Louis, MO). Cultures were run until cell viability dropped below 50%. Supernatant was harvested by pelleting the cells at 250g for 30 minutes followed by pre-filtration through Nalgene™ Glass Pre-filters (Thermo Scientific, Waltham, MA) and 0.45 micron SFCA filtration (Nalgene, Thermo Scientific, Waltham, MA), then stored frozen at –20°C until purification. Proteins were purified using an N-terminal affinity tag derived from type 1 herpes simplex virus glycoprotein D (gD) as previously described (16).

### Glycosidase digestion and SDS-PAGE

Endo H and PNGase F (New England BioLabs, Ipswich, MA) digests were performed per the manufacturer’s protocol on 5ug of purified envelope protein using one unit of glycosidase. Samples were reduced and denatured then digested overnight at 37°C. Digested samples were run on NuPAGE (Thermo Fisher, Invitrogen, Carlsbad, CA) 4–12% BisTris precast gels in MES running buffer then stained with SimplyBlue stain (Thermo Fisher, Invitrogen, Carlsbad, CA).

### Fluorescence immunoassays to measure antibody binding

A fluorescence immunoassay (FIA) was used to measure the binding of polyclonal or monoclonal antibodies to recombinant envelope proteins. For antibody binding to purified proteins, Greiner Fluortrac 600 microtiter plates (Greiner Bio One, Kremsmünster, Austria) were coated with 2ug/mL of purified envelope protein overnight in PBS with shaking. Plates were blocked in PBS + 2.5% BSA (blocking buffer for 90 minutes, then washed four times with PBS containing 0.05% Tween-20 (Sigma). Serial dilutions of monoclonal antibodies were added in a range from 10ug/mL to 0.0001ug/mL, and then incubated at 25°C for 90 minutes with shaking. After incubation and washing, 488 Alexa Fluor conjugated anti-human or anti-murine (Invitrogen, CA) was added at a 1:3000 dilution in PBS + 1% BSA. Plates were incubated for 90 minutes with shaking then washed four times with 0.05% Tween PBS using an automated plate washer. Plates were then imaged in a plate spectrophotometer (Envision System, Perkin Elmer) at excitation and emission wavelengths of 395nm and 490nm respectively. For antibody binding to unpurified envelope proteins in cell culture supernatants, Greiner Fluortrac 600 microtiter plates (Greiner Bio-one, Germany) were coated with 2ug/mL of purified mouse monoclonal antibody to an epitope in the V2 domain (10C10) or the gD purification tag (34.1) overnight in PBS with shaking. Plates were blocked in PBS + 2.5% BSA blocking buffer for 90 minutes, then washed four times with PBS containing 0.05% Tween-20 (Sigma). 150μl of 40x diluted supernatant were then added to each well or 10μg/mL of purified protein in control lanes, then incubated at 25°C for 90minutes with shaking. After incubation and washing, PG9 was added in a range from 10μg/mL to 0.0001μg/mL, and then incubated at 25°C for 90minutes with shaking. After incubation and washing, fluorescently conjugated anti-human or anti-murine (Invitrogen, CA) was added at a 1:3000 dilution. Plates were incubated for 90 minutes with shaking then washed four times with 0.05% Tween PBS using an automated plate washer. Plates were then imaged in a plate spectrophotometer (Envision System, Perkin Elmer, Waltham, MA) at excitation and emission wavelengths of 395nm and 490nm respectively.

The broadly neutralizing monoclonal antibody PG9 was purchased from Polymun (Klosterneuburg, Austria) or produced in-house using 293 HEK cells from a synthetic gene created on the basis of published sequence data (available from the NIH AIDS Reagent Program, Germantown, MD). Alexa Fluor 488 conjugated anti-human IgG, anti-rabbit IgG, and anti-mouse IgG polyclonal antibodies were obtained from Invitrogen (Invitrogen, Thermo Fisher, Carlsbad, CA)

### Glycan composition analysis by MALDI-TOF-MS

Glycoprotein sample (∼100 μg) suspended in 50 mM ammonium bicarbonate buffer was incubated with trypsin (5μg, Sigma Aldrich), for 18 h at 37°C. Digested peptides were desalted and purified by passing through a C18 sep-pak cartridge after inactivating trypsin by heating at 95°C for 5 min. The purified peptides were then treated with PNGase F (23 IUB milliunits, NEB#P0705, New England BioLabs, Ipswich, MA) at 37 ˚C for 16 h, to release the N-glycans. The released N-glycans were desalted and purified from the peptides by C18 sep-pak cartridge followed by freeze-drying. Finally, the N-glycans were permethylated for 10 min at RT, by using 100 μl of methyl iodide in the presence of NaOH/DMSO base (350 μl). The reaction was quenched by adding water (1 ml) and the permethylated N-glycans were extracted by organic phase separation using dichloromethane (2 ml). The excess of dichloromethane was removed by stream of nitrogen and subsequently, prepared for MALDI-MS analysis (84).

Permethylated N-glycans were dissolved in methanol (20 μl) and small aliquot (∼1 μl) was spotted on to MALDI plate (Opti-TOF-384 well insert, Applied Biosystems, Foster City, CA) and crystallized with DHB matrix (20 mg/ml in 50 % Methanol/water, Sigma Aldrich). Data were obtained from AB SCIEX MALDI TOF/TOF 5800 (Applied Biosystem MDS Analytical Technologies, Foster City, CA) mass spectrometer in reflector positive-ion mode. Data analysis was performed by using Data Explorer V4.5, and the assignment of glycan structure was based on the primary mass (m/z) coupled with MS/MS fragmentation profile using the Expasy database online and the glycowork bench software analysis (84, 85).

### Minute Virus of Mice infectivity assay and sterility testing

IDEXX BioResearch (Columbia, MO) performed a Minute Virus of Mice (MVM) infectivity assay. Cells were cultured at 4×10^5^ cells/mL, in 100mL total volume under conditions described above in a spinner flask for five days. CHO-S and MGAT1^-^ CHO cells were infected with 1 multiplicity of infection (MOI) of MVM prototypic strain (MVMp) or MVM Cutter strain (MVMc) and evaluated in triplicate. Five mL aliquots were removed on days 1, 3, and 5, and cells were pelleted by centrifugation and stored at –20°C. Day 5 samples were evaluated by quantitative polymerase chain reaction (qPCR) for MVM and 18S using proprietary primers. The qPCR crossing point (CP) values were reported and copy numbers based upon standard curves.

The cell line was tested for a panel of adventitious agents, cell line species, and in-vitro virus contamination by IDEXX BioResearch (Columbia, MO, USA) using a PCR based protocol described in supplemental materials.

## Acknowledgments

Dr. Benjamin Abrams, UCSC Life Science Microscopy Center, provided technical support. Dr. Matthew H. Myles at IDEXX BioResearch developed and performed the MVM infectivity assays. Chelsea Didinger of the Department of Biomolecular Engineering at UCSC assisted with text and manuscript preparation.

